# A phylodynamic workflow to rapidly gain insights into the dispersal history and dynamics of SARS-CoV-2 lineages

**DOI:** 10.1101/2020.05.05.078758

**Authors:** Simon Dellicour, Keith Durkin, Samuel L. Hong, Bert Vanmechelen, Joan Martí-Carreras, Mandev S. Gill, Cécile Meex, Sébastien Bontems, Emmanuel André, Marius Gilbert, Conor Walker, Nicola De Maio, Nuno R. Faria, James Hadfield, Marie-Pierre Hayette, Vincent Bours, Tony Wawina-Bokalanga, Maria Artesi, Guy Baele, Piet Maes

## Abstract

Since the start of the COVID-19 pandemic, an unprecedented number of genomic sequences of the causative virus (SARS-CoV-2) have been generated and shared with the scientific community. The unparalleled volume of available genetic data presents a unique opportunity to gain real-time insights into the virus transmission during the pandemic, but also a daunting computational hurdle if analysed with gold-standard phylogeographic approaches. We here describe and apply an analytical pipeline that is a compromise between fast and rigorous analytical steps. As a proof of concept, we focus on the Belgium epidemic, with one of the highest spatial density of available SARS-CoV-2 genomes. At the global scale, our analyses confirm the importance of external introduction events in establishing multiple transmission chains in the country. At the country scale, our spatially-explicit phylogeographic analyses highlight that the national lockdown had a relatively low impact on both the lineage dispersal velocity and the long-distance dispersal events within Belgium. Our pipeline has the potential to be quickly applied to other countries or regions, with key benefits in complementing epidemiological analyses in assessing the impact of intervention measures or their progressive easement.

---

First reported in early December 2019 in the province of Hubei (China), COVID-19 (coronavirus disease 2019) is caused by a new coronavirus (severe acute respiratory syndrome coronavirus 2; SARS-CoV-2) that has since rapidly spread around the world^1,2^, causing an enormous public health and social-economic impact^3,4^. Since the early days of the pandemic, there has been an important mobilisation of the scientific community to understand its epidemiology and help providing a real-time response. To this end, research teams around the world have massively sequenced and publicly released dozens of thousands of viral genome sequences to study the origin of the virus^5,6^, and to trace its spread at global, country or community-level scales^7–9^. In this context, a platform like Nexstrain^10^, already widely used and recognised by the academic community and beyond, has quickly become a reference to follow the travel history of SARS-CoV-2 lineages.

In the context of the COVID-19 pandemic, the volume of genomic data available presents a unique opportunity to gain valuable real-time insights into the dispersal dynamics of the virus. Yet, the number of available viral genomes is increasing every day, leading to substantial computational challenges. While Bayesian phylogeographic inference represents the gold standard for inferring the dispersal history of viral lineages^11^, these methods are computationally intensive and will fail to provide useful results in an acceptable amount of time.

To tackle this practical limitation, we here describe and apply an analytical pipeline that is a compromise between fast and rigorous analytical steps. In practice, we propose to take advantage of the rapid time-scaled phylogenetic tree inference process used by the online Nextstrain platform^10^. Specifically, we aim to use the resulting time-scaled tree as a fixed empirical tree along which we infer the ancestral locations with the discrete^12^ and spatially-explicit^13^ phylogeographic models implemented in the software package BEAST 1.10^14^.

In Belgium, there are two main different laboratories (from the University of Leuven and the University of Liège) involved in sequencing SARS-CoV-2 genomes extracted from confirmed COVID-19 positive patients. To date, some genomes (*n*=58) have also been sequenced at the University of Ghent, but for which metadata about the geographic origin are unavailable. As of June 10, 2020, a total of 740 genomes have been sequenced by these research teams and deposited on the GISAID (Global Initiative on Sharing All Influenza Data^15^) database. In the present study, we exploit this comprehensive data set to unravel the dispersal history and dynamics of SARS-CoV-2 viral lineages in Belgium. In particular, our objective is to investigate the evolution of the circulation dynamics through time and assess the impact of lockdown measures on spatial transmission. Specifically, we aim to use phylogeographic approaches to look at the Belgian epidemic at two different levels: (i) the importance of introduction events into the country, and (ii) viral lineages circulating at the nationwide level.

## RESULTS

### Importance of introduction events into the country

On June 10, 2020, we downloaded all Belgian SARS-CoV-2 sequences (*n*=740) available on GISAID, as well as non-Belgian sequences (4,309) originated from 126 different countries and used in Nextstrain to represent the overall dispersal history of the virus. We generated a time-scaled phylogenetic tree using a rapid maximum likelihood approach^16^ and subsequently ran a preliminary discrete phylogeographic analysis along this tree to identify internal nodes and descending clades that likely correspond to distinct introductions into the Belgian territory (Fig. 1, S2). We inferred a minimum number of 331 introduction events (95% HPD interval = [315-344]). When compared to the number of sequences sampled in Belgium (*n*=740), this number illustrates the relative importance of external introductions in establishing transmission chains in the country. Introduction events resulted in distinct clades (or “clusters”) linking varying numbers of sampled sequences (Fig. 1). However, many clusters only consisted of one sampled sequence. According to the time-scaled phylogenetic tree and discrete phylogeographic reconstruction (Fig. S1), some of these introduction events could have occurred before the return of Belgian residents from carnival holidays (around March 1, 2020), which was considered as the major entry point of transmission chains in Belgium.

**Figure 1.**
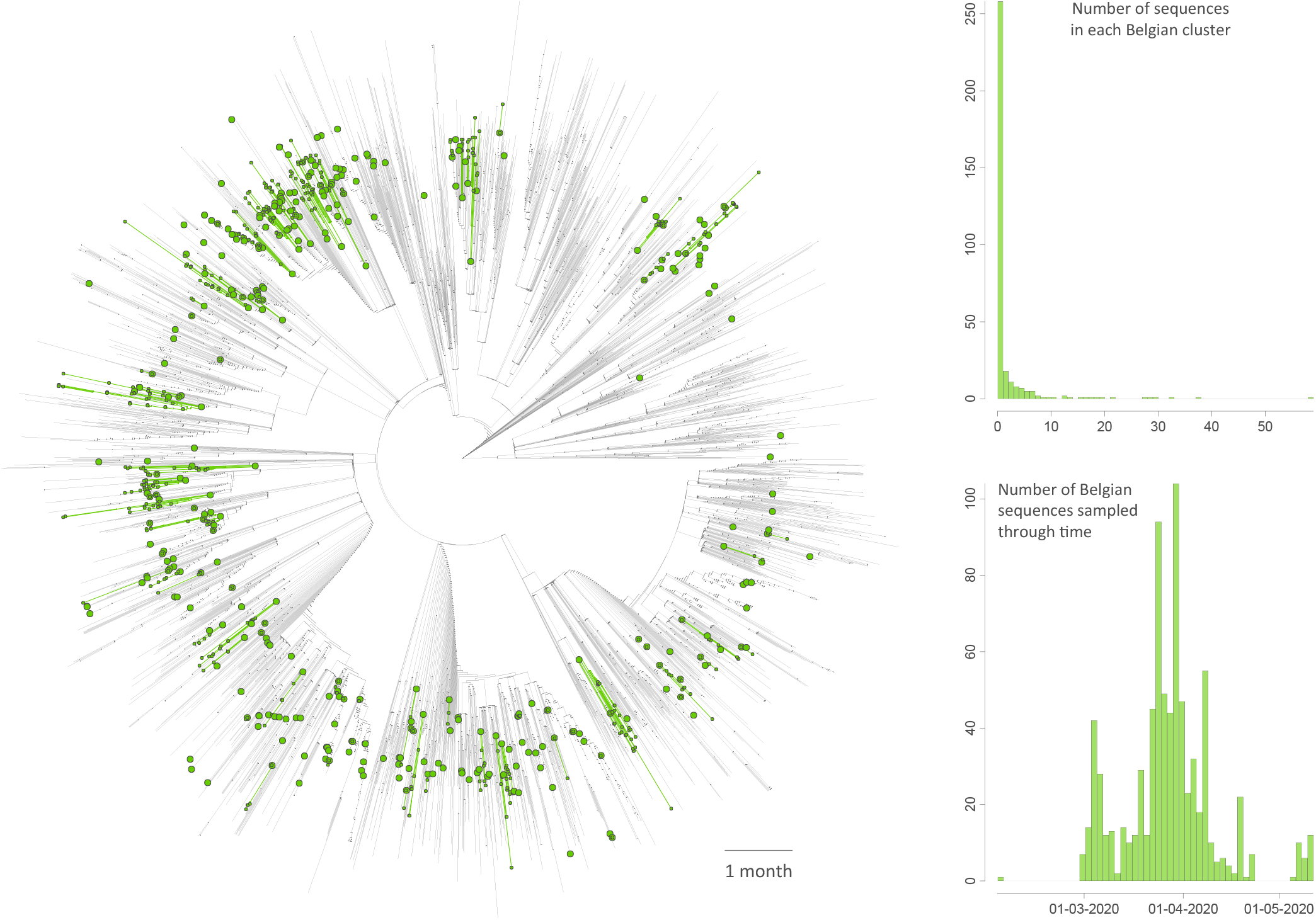
Time-scaled phylogenetic tree in which we identified Belgian clusters. A cluster is here defined as a phylogenetic clade likely corresponding to a distinct introduction into the study area (Belgium). We delineated these clusters by performing a simplistic discrete phylogeographic reconstruction along the time-scaled phylogenetic tree while only considering two potential ancestral locations: “Belgium” and “non-Belgium”. We identified a minimum number of 331 lineage introductions (95% HPD interval = [315-344]), which gives the relative importance of external introductions considering the number of sequences currently sampled in Belgium (740). On the tree, lineages circulating in Belgium are highlighted in green, and green nodes correspond to the most ancestral node of each Belgian cluster (see also Figure S1 for a non-circular visualisation of the same tree). Besides the tree, we also report the distribution of cluster sizes (number of sampled sequences in each cluster) as well as the number of sequences sampled through time.

### Impact of lockdown measures at the country level

To analyse the circulation dynamics of viral lineages within the country, we then performed spatially-explicit phylogeographic inference along the previously identified Belgian clades (Fig. 2A). Our reconstructions reveal the occurrence of long-distance dispersal events both before (Fig. 2B) and during (Fig. 2C) the lockdown. By placing phylogenetic branches in a geographical context, spatially-explicit phylogeographic inference allows treating those branches as conditionally independent movement vectors^17^. Here, we looked at these movement vectors to assess how the dispersal dynamics of lineages was impacted by the national lockdown, of which the main measures were implemented on March 18, 2020.

**Figure 2.**
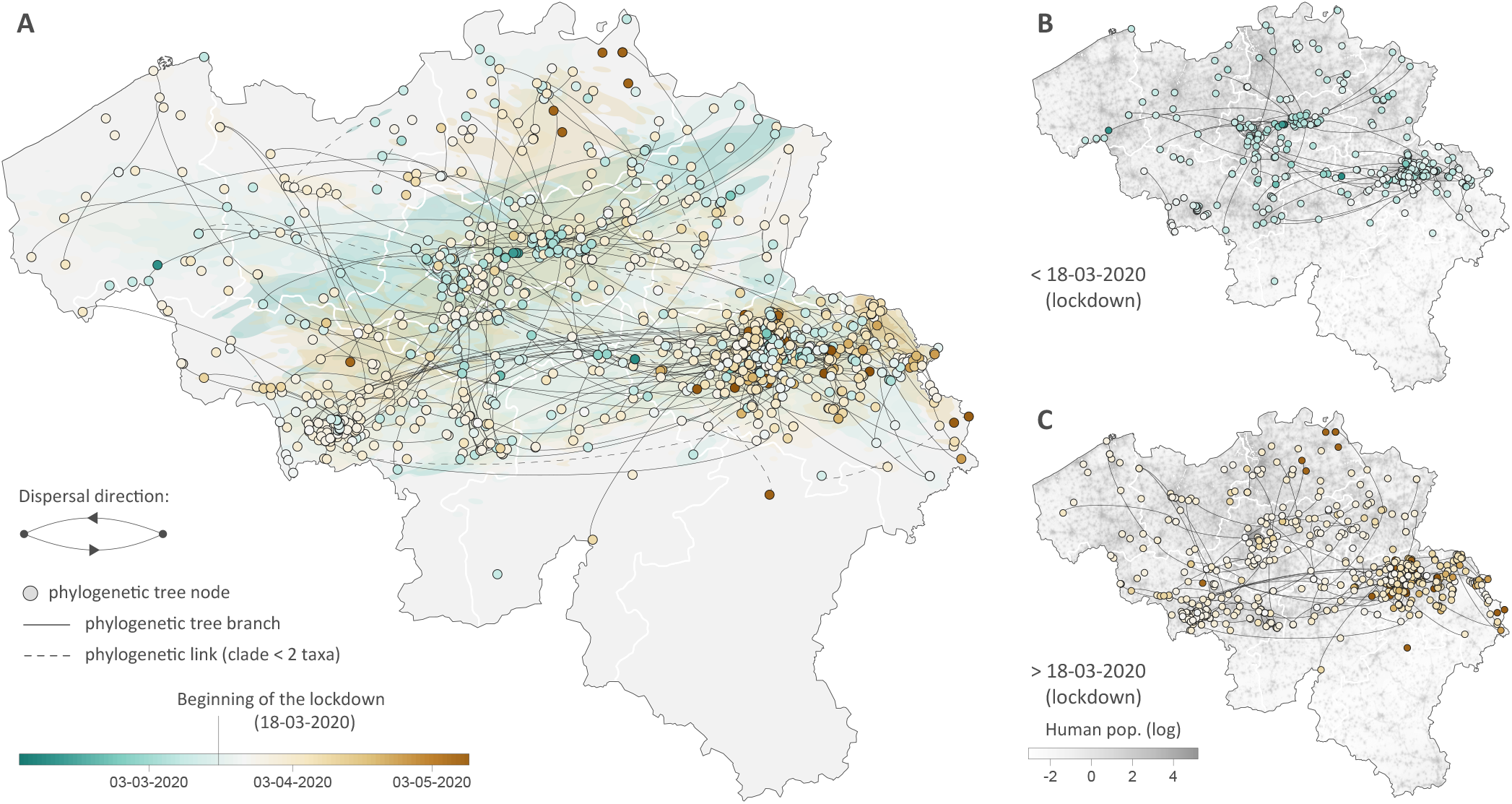
Spatially-explicit phylogeographic reconstruction of the dispersal history of SARS-CoV-2 lineages in Belgium. (**A**) Continuous phylogeo-graphic reconstruction performed along each Belgian clade (cluster) identified by the initial discrete phylogeographic analysis. For each clade, we mapped the maximum clade credibility (MCC) tree and overall 80% highest posterior density (HPD) regions reflecting the uncertainty related to the phylogeographic inference. MCC trees and 80% HPD regions are based on 1,000 trees subsampled from each post burn-in posterior distribution. MCC tree nodes were coloured according to their time of occurrence, and 80% HPD regions were computed for successive time layers and then superimposed using the same colour scale reflecting time. Continuous phylogeographic reconstructions were only performed along Belgian clades linking at least three sampled sequences for which the geographic origin was known (see the Methods section for further details). Besides the phylogenetic branches of MCC trees obtained by continuous phylogeographic inference, we also mapped sampled sequences belonging to clades linking less than three geo-referenced sequences. Furthermore, when a clade only gathers two geo-referenced sequences, we highlighted the phylogenetic link between these two sequences with a dashed curve connecting them. Sub-national province borders are represented by white lines. (**B**) MCC tree branches occurring before the 18^th^ March 2020 (beginning of the lockdown). (**C**) MCC tree branches occurring after the 18^th^ March 2020. See also Figure S2 for a zoomed version of the dispersal history of viral lineages in the Province of Liège, for which we have a particularly dense sampling.

Firstly, we investigated if the lockdown was associated with a change in lineage dispersal velocity. We estimated a substantially higher dispersal velocity before the lockdown (5.4 km/day, 95% HPD [5.0-6.1]) compared to during the lockdown (2.4 km/day, 95% HPD [2.3-2.5]). This trend is further confirmed when focusing on the Province of Liège for which we have a particularly dense sampling: in that province, we estimated a lineage dispersal velocity of 2.8 km/day (95% HPD [2.2-3.6]) before the lockdown and of 1.1 km/day (95% HPD [1.0-1.2]) during the lockdown. However, the evolution of the dispersal velocity through time is less straightforward to interpret (Fig. 3): while the lineage dispersal velocity was globally higher at the early phase of the Belgian epidemic, which corresponds to the week following the returns from carnival holidays, it then seemed to drop just before the beginning of the lockdown before increasing again to reach a plateau. In the second half of April, our estimates indicate that the lineage dispersal velocity drops again. However, this result may be an artefact associated with the notably lower number of phylogenetic branches currently inferred during that period (Fig. 3).

**Figure 3.**
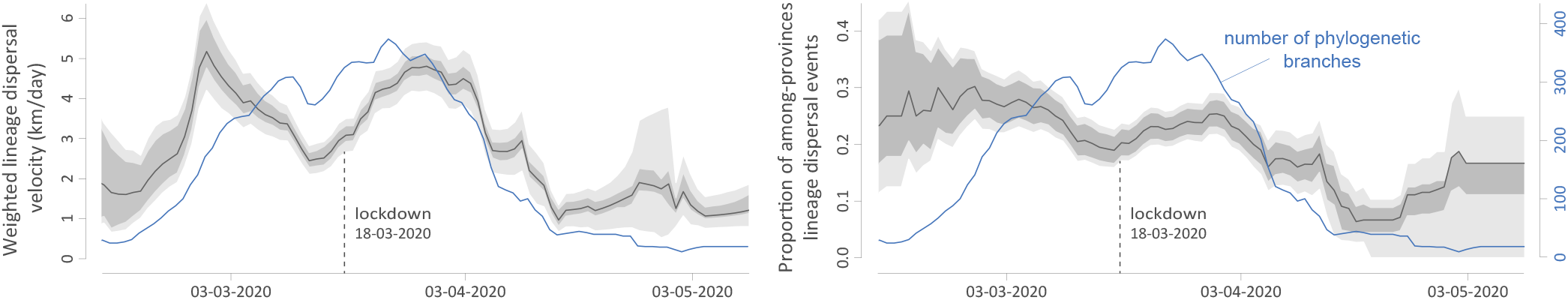
Evolution of viral lineage dispersal dynamics during the Belgian epidemic. These estimates are based on 1,000 trees subsampled from each post burn-in posterior distribution. Except the number of phylogenetic branches occurring at each time slice, all estimates were smoothed using a 14-days sliding window. Dark grey surrounding polygons represent 95% credible intervals, and light grey surrounding polygons represent 95% credible intervals re-estimated after subsampling 75% of branches in each of the 1,000 posterior trees. The credible interval based on the subsampling procedure is an indication of the robustness of the estimate. In addition, we also report the number of phylogenetic branches occurring per tree at each time slice (blue curve). The number of branches available at each time slice is an additional, yet qualitative, indication of robustness of the estimate for a given time period.

Secondly, we further investigated the impact of the lockdown on the dispersal events among provinces. Our analyses indicate that among-provinces dispersal events tended to decrease during the epidemic (Fig. 3): such dispersal events were more frequent at the beginning of the epidemic and then progressively decreased until reaching a plateau at the beginning of the lockdown. Again, the relatively limited number of phylogenetic branches currently inferred from mid-April does not really allow to interpret the fluctuations of the proportion of among-provinces dispersal events during that period.

## DISCUSSION

Our preliminary phylogeographic investigation reveals the important contribution of external introduction events for the establishment of the SARS-CoV-2 epidemic in Belgium. This highlights that transmission chains circulating in Belgium were not established by a relatively restricted number of isolated infectious cases, e.g. people returning from skiing holidays in northern Italy. On the contrary, we identify a large number of distinct clades given the number of analysed sequences sampled in Belgium. This overall observation is in line with other reports, e.g. in California where no predominant lineage was identified either^18^.

Our spatially-explicit phylogeographic analyses uncover the spatiotemporal distribution of Belgian SARS-CoV-2 clusters, indicating a relatively low impact of the lockdown on both the dispersal velocity of viral lineages and on the frequency of long-distance dispersal events. While it has been demonstrated that the national lockdown had an overall impact on the virus transmission, i.e. reducing its effective reproduction number to a value below one^19^, our results highlight that the lockdown did not clearly decrease the velocity at which the viral lineages travelled or their ability to disperse over long distances within the country. This finding may be important to consider in the context of potential future lockdown measures, especially if more localised (e.g. at the province or city level). Indeed, locally reduced transmission rates will not automatically be associated with a notable decrease in the average velocity or distance travelled by lineage dispersal events, which could in turn limit the effectiveness of localised lockdown measures in containing local upsurge of the virus circulation.

Applying the present phylodynamic pipeline in a real-time perspective does not come without risk as new sequences can sometimes be associated with spurious nucleotide changes that could be associated with sequencing or assembling errors. Directly starting from inference results kept up to date by a database like GISAID allows for fast analytical processing but also relies on newly deposited data that could sometimes carry potential errors. To remedy such potentially challenging situations, our proposed pipeline could be extended with a sequence data resource component that makes uses of expert knowledge regarding a particular virus. The GLUE^20^ software package allows new sequences to be systematically checked for potential issues, and could hence be an efficient tool to safely work with frequently updated SARS-CoV-2 sequencing data. Such a “CoV-GLUE” resource is currently being developed (http://cov-glue.cvr.gla.ac.uk/#/home).

While we acknowledge that a fully integrated analysis (i.e. an analysis where the phylogenetic tree and ancestral locations are jointly inferred) would be preferable, fixing an empirical time-scaled tree represents a good compromise to rapidly gain insights into the dispersal history and dynamics of SARS-CoV-2 lineages. Indeed, the number of genomes available, as well as the number of different sampling locations to consider in the analysis, would lead to a joint analysis requiring weeks of run-time in a Bayesian phylogenetic software package like BEAST. To illustrate the computational demands of such approach, we ran a classic Bayesian phylogenetic analysis on a smaller SARS-CoV-2 data set (2,795 genomic sequences) using BEAST 1.10 (data not shown). This analysis required over 150 hours to obtain enough samples from the joint posterior, while using the latest GPU accelerated implementations^21^ on 15 parallel runs. With a combined chain length of over 2.2×10^9^ states, and an average runtime of 0.9 hours per million states, the significant computational demands required make this approach impractical when speed is critical. On the other hand, we here use a maximum likelihood method implemented in the program TreeTime^16^ to infer a time-scaled phylogenetic tree in a short amount of time (~3 hours for the data set analysed here). Given the present urgent situation, we have deliberately assumed a time-scaled maximum-likelihood phylogenetic tree as a fair estimate of the true time-scaled phylogenetic tree.

Our analytical workflow has the potential to be rapidly applied to study the dispersal history and dynamics of SARS-CoV-2 lineages in other restricted or even much broader study areas. We believe that spatially-explicit reconstruction can be a valuable tool for highlighting specific patterns related to the circulation of the virus or assessing the impact of intervention measures. While new viral genomes are sequenced and released daily, a limitation could paradoxically arise from the non-accessibility of associated metadata. Indeed, without sufficiently precise data about the geographic origin of each genome, it is not feasible to perform a spatially-explicit phylogeographic inference. In the same way that viral genomes are deposited in databases like GISAID, metadata should also be made available to enable comprehensive epidemiological investigations with a similar approach as we presented here.

## METHODS

### SARS-CoV-2 sequencing in Belgium

At the University of Leuven, RNA extracts from SARS-CoV-2 infected patients were selected anonymously and based on a city?s postal code within Belgium. These RNA extracts were provided by the National Reference Center for Coronaviruses and UZ Leuven. Reverse transcription was carried out via SuperScript IV and cDNA was posteriorly amplified using Q5^®^ High-Fidelity DNA Polymerase (NEB) with the ARTIC nCov-2019 primers and following the recommendations in the sequencing protocol of the ARTIC Network (https://artic.network/ncov-2019). Samples were multiplexed following the manufacturer’s recommendations using the Oxford Nanopore Native Barcoding Expansion kits NBD104 (1-12) and NBD114 (13-24), in conjunction with Ligation Sequencing Kit 109 (Oxford Nanopore). Sequencing was carried out on a MinION sequencer using R9.4.1 flow cells and MinKNOW 2.0 software.

At the University of Liège, RNA was extracted from clinical samples (300μl) via a Maxwell 48 device using the Maxwell RSC Viral TNA kit (Promega) with a viral inactivation step using Proteinase K, following the manufacturer’s instructions. RNA elution occurred in 50μl of RNAse free water. Reverse transcription was carried out via SuperScript IV VILOTM Master Mix, and 3.3μl of the eluted RNA was combined with 1.2μl of master mix and 1.5μl of H_2_O. This was incubated at 25°C for 10 min, 50°C for 10 min and 85°C for 5 min. PCRused Q5^®^ High-Fidelity DNA Polymerase (NEB), the primers and conditions followed the recommendations in the sequencing protocol of the ARTIC Network. Samples were multiplexed following the manufacturer’s recommendations using the Oxford Nanopore Native Barcoding Expansion kits 1-12 and 13-24, in conjunction with Ligation Sequencing Kit 109 (Oxford Nanopore). Sequencing was carried out on a Minion using R9.4.1 flow cells. Data analysis followed the SARS-CoV-2 bioinformatics protocol of the ARTIC Network.

### Inference of a time-scaled phylogenetic tree

To infer our time-scaled phylogenetic tree, we selected all non-Belgian sequences in the Nextstrain analysis, along with all available Belgian sequences in GISAID to be included in our analysis as of June 10, 2020. Once we knew which were the accessions of interest, we downloaded the latest whole genome alignment from GISAID and removed all non-relevant sequences. We then cleaned the alignment by manually trimming the 5’ and 3’ untranslated regions (RefSeq NC_045512.2) and gap-only sites. To obtain a maximum-likelihood phylogeny, we ran IQ-TREE 2.0.3^22^ under a general time reversible^23^ (GTR) model of nucleotide substitution with empirical base frequencies and four Free Rate^24^ site categories. This model configuration was selected as the best GTR model using IQ-TREE?s ModelFinder tool. The tree was then inspected for outlier sequences using TempEst^25^ 1.5.3 and, once the outliers were removed, time-calibrated using TreeTime 0.7.420. To replicate the Nextstrain workflow as closely as possible, we specified a clock rate of 8×10^-4^ in TreeTime and removed samples that deviate more than four interquartile ranges from the root-to-tip regression.

### Preliminary discrete phylogeographic analysis

We performed a preliminary phylogeographic analysis using the discrete diffusion model^12^ implemented in the software package BEAST 1.10^14^. The objective of this first analysis was to identify independent introduction events of SARS-CoV-2 lineages into Belgium. To this end, we used our time-scaled phylogenetic tree as a fixed empirical tree and only considered two possible ancestral locations: *Belgium* and *non-Belgium*. Bayesian inference through Markov chain Monte Carlo (MCMC) was run on this empirical tree for 10^6^ generations and sampled every 1,000 generations. MCMC convergence and mixing properties were inspected using the program Tracer 1.7^26^ to ensure that effective sample size (ESS) values associated with estimated parameters were all >200. After having discarded 10% of sampled trees as burn-in, a maximum clade credibility (MCC) tree was obtained using TreeAnnotator 1.10^14^. We used the resulting MCC tree to delineate Belgian clusters here defined as phylogenetic clades corresponding to independent introduction events in Belgium.

### Continuous and post hoc phylogeographic analyses

We used the continuous diffusion model^13^ available in BEAST 1.10^14^ to perform a spatially-explicit (or “continuous”) phylogeographic reconstruction of the dispersal history of SARS-CoV-2 lineages in Belgium. We employed a relaxed random walk (RRW) diffusion model to generate a posterior distribution of trees whose internal nodes are associated with geographic coordinates^13^. Specifically, we used a Cauchy distribution to model the among-branch heterogeneity in diffusion velocity. We performed a distinct continuous phylogeographic reconstruction for each Belgian clade identified by the initial discrete phylogeographic inference, again fixing a time-scaled subtree as an empirical tree. As phylogeographic inference under the continuous diffusion model does not allow identical sampling coordinates assigned to the tips of the tree, we avoided assigning sampling coordinates using the centroid point of each administrative area of origin. For a given sampled sequence, we instead retrieved geographic coordinates from a point randomly sampled within its municipality of origin, which is the maximal level of spatial precision in available metadata. This approach avoids using the common “jitter” option that adds a restricted amount of noise to duplicated sampling coordinates. Using such a jitter could be problematic because it can move sampling coordinates to administrative areas neighbouring their actual administrative area of origin^27^. Furthermore, the administrative areas considered here are municipalities and are rather small (there are currently 581 municipalities in Belgium). The clade-specific continuous phylogeographic reconstructions were only based on Belgian tip nodes for which the municipality of origin was known, i.e. 639 out of 740 genomic sequences. Furthermore, we only performed a continuous phylogeographic inference for Belgian clades linking a minimum of three tip nodes with a known sampling location (municipality).

Each Markov chain was run for 10^6^ generations and sampled every 1,000 generations. As with the discrete phylogeographic inference, MCMC convergence/mixing properties were assessed with Tracer, and MCC trees (one per clade) were obtained with TreeAnnotator after discarding 10% of sampled trees as burn-in. We then used functions available in the R package “seraphim”^28,29^ to extract spatiotemporal information embedded within the same 1,000 posterior trees and visualise the continuous phylogeographic reconstructions. We also used “seraphim” to estimate the following weighted lineage dispersal velocity, and we verified the robustness of our estimates through a subsampling procedure consisting of re-computing the weighted dispersal velocity after having randomly discarded 25% of branches in each of the 1,000 posterior trees. The weighted lineage dispersal velocity is defined as follows, where *d_i_* and *t_i_* are the geographic distance travelled (great-circle distance in km) and the time elapsed (in days) on each phylogeny branch, respectively:

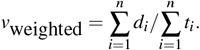

## Data accessibility

The new sequences have been deposited in GISAID and all data (sequence metadata, BEAST input, output files, and R scripts for our analyses) are available in the following GitHub repository: https://github.com/sdellicour/sars_cov_2_pipeline.

## Acknowledgments

We are grateful to Sébastien Kozlowskyj for his assistance in using the Dragon2 cluster of the University of Mons, and to Ine Boonen for her assistance during SARS-CoV-2 sequencing. SD and MG are supported by the Fonds National de la Recherche Scientifique (FNRS, Belgium). KD and MA, and the ULiège sequencing effort, are supported by the grant WALGEMED from the Walloon Region (convention n°1710180). BV is supported by a FWO SB grant for strategic basic research of the “Fonds Wetenschappelijk Onderzoek”/Research foundation Flanders (1S28617N). JMC is supported by a doctoral grant from HONOURs (Host switching pathogens, infectious outbreaks and zoonosis) Marie-Sklodowska-Curie training network (721367). NRF is supported by a Sir Henry Dale Fellowship (204311/Z/16/Z) and by a MRC/FAPESP CADDE partnership award (MR/S0195/1). TWB is supported by the Special Research Fund, KU Leuven (Bijzonder Onderzoeksfonds, KU Leuven, 3M170314 C14/17/100). GB acknowledges support from the Interne Fondsen KU Leuven / Internal Funds KU Leuven under grant agreement C14/18/094, and the Research Foundation Flanders (“Fonds voor Wetenschappelijk Onderzoek Vlaanderen”, G0E1420N). Computational resources have been provided by the Consortium des Équipements de Calcul Intensif (CÉCI), funded by the Fonds de la Recherche Scientifique de Belgique (FNRS) under Grant No. 2.5020.11 and by the Walloon Region. We are grateful to all of the research teams that have deposited SARS-CoV-2 genome data on GISAID.

## Notes

https://github.com/sdellicour/covid19_spell

### Competing Interest Statement

The authors have declared no competing interest.

