## Supplementary material for "A phylodynamic workflow to rapidly gain insights into the dispersal history and dynamics of SARS-CoV-2 lineages": Figures S1-S2

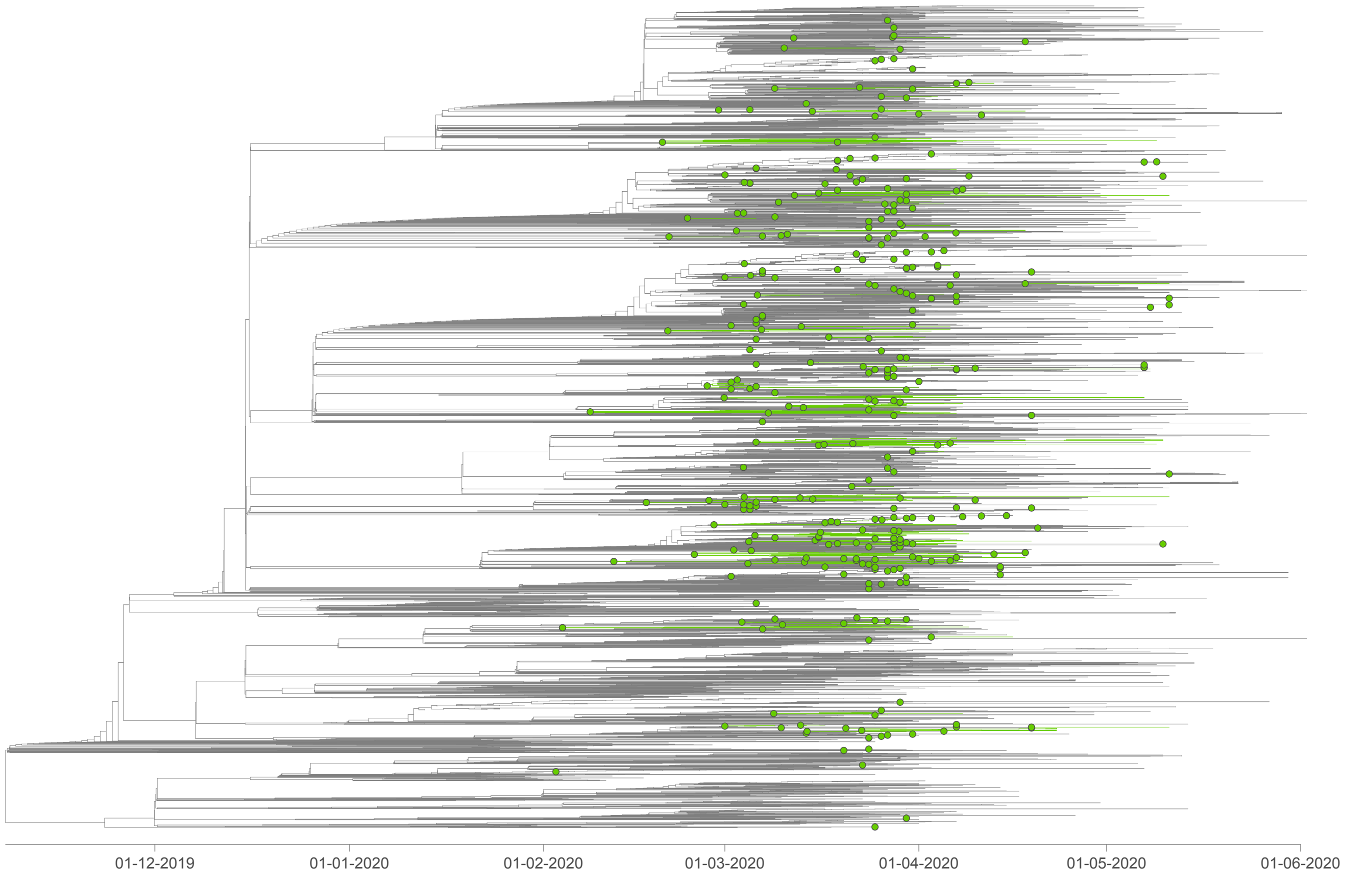


**Figure S1. Time-scaled phylogenetic tree in which we identified Belgian clusters.** A cluster is here defined as a phylogenetic clade likely corresponding to a distinct introduction into the Belgian territory. We delineated these clusters by performing a simplistic discrete phylogeographic reconstruction along the time-scaled phylogenetic tree while only considering two potential ancestral locations: "Belgium" and "outside Belgium". On the tree, lineages circulating in Belgium are highlighted in green, and green nodes correspond to the most ancestral node of each Belgian cluster.

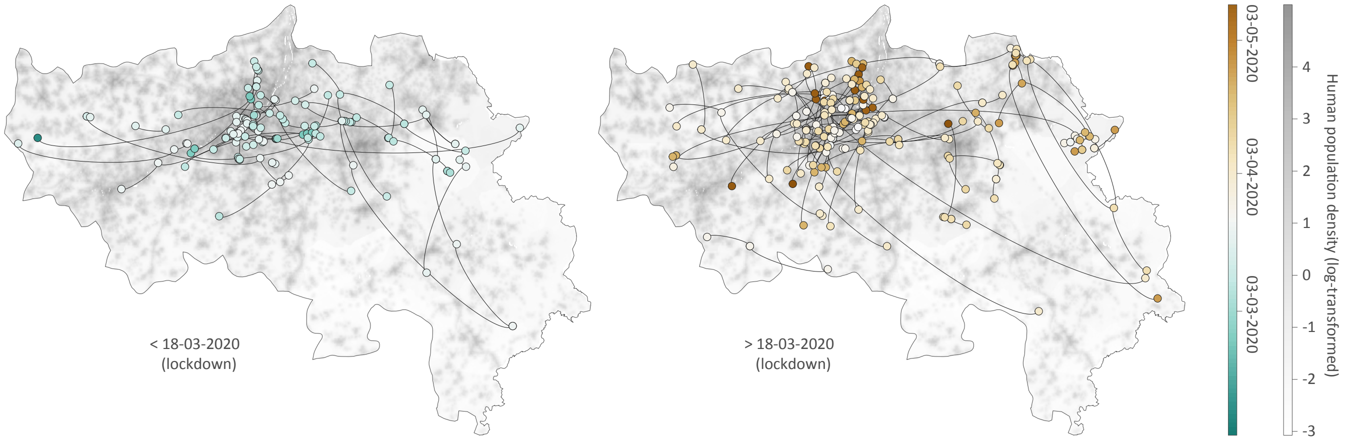


**Figure S2. Spatially-explicit phylogeographic reconstruction of the dispersal history of SARS-CoV-2 lineages in the Province of Liège.** Continuous phylogeographic reconstruction was performed along each Belgian clade (cluster) identified by the initial discrete phylogeographic analysis. For each clade, we mapped the maximum clade credibility (MCC) branches located in the province of Liège, before and after the 18th March 2020 (i.e. the beginning of the lockdown).
